# Identification of lumbrokinase by fibrin-plate method, fibrin-zymography and liquid chromatography mass spectrometry

**DOI:** 10.1101/656827

**Authors:** Bingbing Ke’, Ruiqing Xian, Hong Jiang, Jianghong Guo

**Affiliations:** Hubei institute for drug control, Wuhan China; Shandong institute for food and drug control, Jinan China

**Keywords:** Lumbrokinase, *Eisenia fetida*, *fibrin-zymography*, *liquid chromatography mass spectrometry*

## Abstract

Lumbrokinase, extracted from the cultured earthworm of *Eisenia fetida*, has been widely used as biochemical medicine in China to prevent or treat thrombosis. In the present study, the mechanism of lumbrokinase was investigated using the fibrin plate method. The results revealed that lumbrokinase contained both fibrinolytic and kinase components. The method of fibrin-zymography was used to show the existence and activity of lumbrokinase, and we proved that the fibrin-zymogram gel could be adopted as the identification method of earthworm. Subsequently, the components were identified by mass spectrometry. According to the results, fibrinolytic related components existed in the drug. These proteins were further compared with other serine proteins. The result showed that the identified proteins were similar to human trypsin and bovine trypsin. Besides, some also exhibited similar characteristics with human plasminogen activators. The mentioned results demonstrated that lumbrokinase products contained two major groups of protein components, suggesting two different functions.

## Introduction

Lumbrokinase(LK), which is ubiquitous in earthworms, has been used as traditional medicines for thousands of years in China. It has been manufactured into enteric capsules or enteric-coated tablets since 1990s for its inconvenience for dosing and unpleasant smell. The most common side effect is the hypersensitivity caused by heterogonous proteins. [1–9]

In China, there are five manufacturers producing lumbrokinase drug substances and enteric capsules. The substances are extracted from the cultured earthworm *(Eisenia fetida).* In general, the extraction process of lumbrokinase consists of three major steps. First, the bodies of earthworm are milled and centrifuged to produce the supernatant. Then, the supernatant was purified with DEAE(diethylaminoethyl) column and then powdered though the process of filtration, ultrafiltration and lyophilization. However, the products, due to the rough manufacturing process, contain many different earthworm fibrinolytic enzymes and other contaminants, which may cause side effects. Besides, though it has been used for a long time and considered effective, there is little knowledge about which types of proteins are vital for the drug. Thus, protein components in lumbrokinase need identification.

This study aimed to illustrate the mechanism of lumbrokinase in the production to identify the proteins in lumbrokinase drug substances by ESI-MS(electrospray ionization mass spectrometry) and to provide data support for the enhancement of its quality specification. In this study, we proved that lumbrokinase had two action mechanisms and identified the proteins existed in the lumbrokinase drug substances. Also, the fibrin-zymography technique was recommended as the identification method of earthworms.

## Materials and method

### Materials

Sample loading buffer(5×, no DTT) and electrophoresis buffer(5×) were purchased from Beyotime biotechnology (China). Coomassie brilliant blue, dithiothreitol, formic acid, and PMSF were available from Sigma(USA); Chymotrypsin (Sequencer Grade V106A) was purchased from Promega (USA). Fibrinogen, lumbrokinase standard, thrombin and plasminogen were provided by National Institute for Food and Drug Control (NIFDC, China). Other chemicals were of analytical grades.

### Enzymatic activity research

Sample preparation: For fibrin plate tests, the drugs from five manufacturers dissolved in the 0.9% sodium chloride, and then they were diluted into a range of concentration (2000U/mL,4000U/mL,6000U/mL,8000U/mL and 10000U/mL). The standard of lumbrokinase was diluted to achieve the same concentration.

Preparation of fibrin plate: Fibrin plates were prepared using the standard method. [10] 39mL of the fibrinogen solution(1.5mg/mL, diluted with PBS buffer) was added into the agarose solution(55°C) and mixed toughly. Subsequently, thrombin solution 3.0mL was added, and the solution was transferred to the plastic culture plates. After evenly mixing the solution, the solution stood still for 1 h at ambient temperature until it was solidified. The preparation of fibrin and kinase plates was the same except that 1mLplasminogen solution (3U/mL) was added into the plates.

For activity analysis, 10 μL test solution of different concentrations into the pole was precisely measured, and then the plates were incubated at 37°C for 18h. The diameter of the hydrolyzed clear zone was measured, and the final values were an average of three replicates.

### Fibrin-zymography

Lumbrokinase drug substances dissolved into 3mg/mL with the solution of 0.9% sodium chloride. SDS-fibrin gel was prepared as described by Kim and Choi.[11, 12] The composition of the gel(10%) is listed in Table 1. 10μL of the supernatant from the drug substances which was diluted in the sample loading buffer were loaded into the fibrin gel. After running the gel(80V 30min;120V 60min), it was soaked in the 2.5% Triton-100 solution for 30min and then incubated in the reaction buffer solution (0.05M KH_2_PO_4_,0.04MNaOH;pH7.4) at 37°C for 40 min. The gel was stained with Coomassie Brilliant blue for 2h and then distained. The digested bands reflecting the enzymatic activities were visualized as clear bands of fibrinolysis against a dark-blue background of undigested fibrin substrate.

**Table 1.** Composition of SDS-fibrin polyacrylamide gel

| Components | 10%(Separating gel) | 5%(Concentrated gel) |
| --- | --- | --- |
| Distilled water | 2.0ml | 1.6ml |
| 1.5M Tris-HCl (pH8.8) | 1.25ml | / |
| 1.0M Tris-HCl (pH6.8) | / | 1.0ml |
| 30%acrylamide | 1.7ml | 0.38ml |
| Bovine fibrinogen(25mg/ml) | 0.1ml | / |
| Bovine thrombin(42U/ml) | 0.1ml | / |
| SDS(10%) | 0.05ml | 0.03ml |
| APS(10%) | 0.05ml | 0.03ml |
| TEMED | 0.01ml | 0.008ml |

### Liquid chromatography and mass spectrometry

Enzyme digestion: A solution containing about 1 mg of lumbrokinase per 1 ml was diluted with a solution containing 6 M urea, 50 mM Tris-HCl (pH 8.0) and 5 mM DTT, incubated for 20 min at 95°C, and then cooled to ambient temperature. Subsequently, a 6-fold volume of 50 mM ammonium bicarbonate solution (pH 7.8) was added. Chymotrypsin (chymotrypsin: sample = 1: 200) was added and digested at 37°C for 12h, and then the solution was centrifugated(8000g) for 30 min before loading the sample to the mass spectrometry.

LC-ESI-MS/MS analysis: An amount equivalent to 2μL of the digested peptide underwent LC-MS/MS using a UPLC system and a Q-TOF mass spectrometry(Waters UPLC^®^ G_2_S Q-TOF). Mobile phase A was 0.1% formic acid, while mobile phase B was acetonitrile. The flow rate was 0.3ml/min. The gradient elution program was allowed for 5min at 5%B, followed by a 40min step that raised eluent B to 40%, a 10 min washing at 40%, and equilibration at 5%B. The total analysis time was 60min. For mass spectrometry, the parameters included: capillary voltage of 3KV, first cone voltage of 36V, sample cone voltage of 60V, source offset voltage of 80V, source temperature at 100 °C, desolvation temperature at 400 °C, cone hole gas 50 L / h, dissolvent gas flow rate of 800 L/h, and the acquisition range : 50 m/z-2000 m/z. The collected data were searched by ProteinLynxGlobal Server3.0 software, Database of *Eisenia foetida* was downloaded from the uniprot website(http://www.uniprot.org/). The protein identification was performed using the following searching parameters. Enzyme: trypsin; Protein mass range:10-150KD; Tolerance: 50ppm; Missedcleavage:1; Fixed modification: carbamidomethylation; Variable modification: methionine oxidation.

## Results

### Enzyme activity measurement

The fibrin plate was used to ascertain the direct fibrinolytic activity of lumbrokinase, as shown in Fig 1a. Besides, the fibrin-plasminogen plate was used to measure the fibrinolytic and kinase activity of lumbrokinase, as shown in Fig 1b. The results revealed that the diameter of the fibrin-plasminogen plate was significantly larger than that of the fibrin plate (Table 2), indicating that lumbrokinase not only exhibited the direct degradation activity of fibrin, but also the ability of activating plasminogen to produce indirect fibrinolysis, showing a kinase activity.

**Fig 1.**
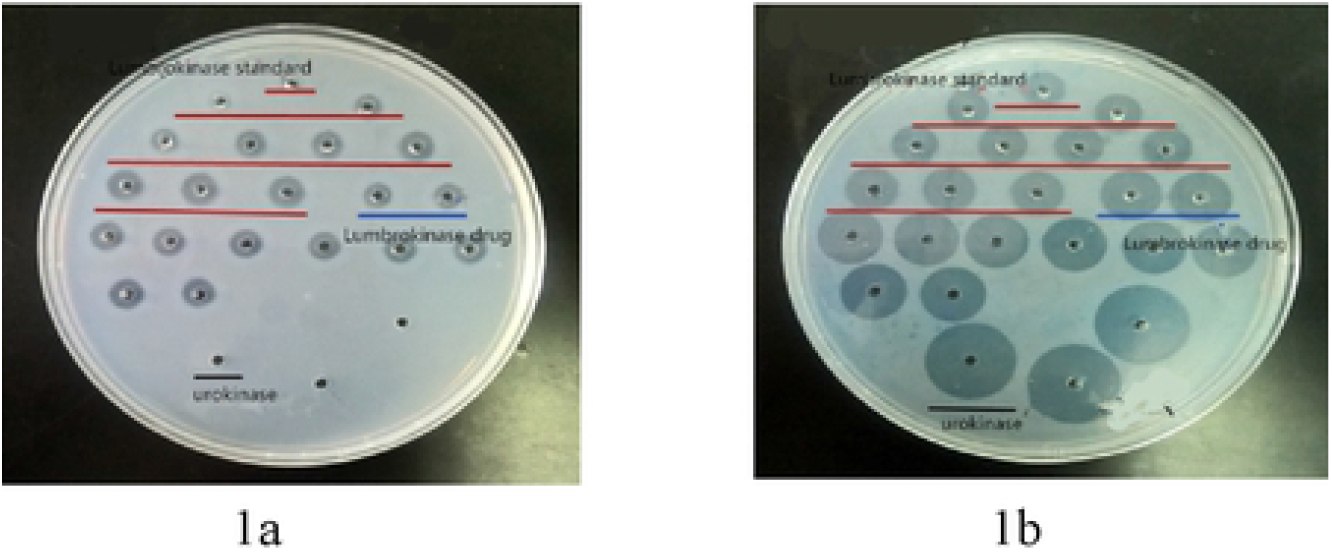
Diameter on the fibrin plate(1a)and the fibrin-plasminogen plate(1b)

**Table 2.** Diameter (mm) on the fibrin plate and the fibrin-plasminogen plate between sample and standard

| Sample concentration(U/mL) |  | 2000 | 4000 | 6000 | 8000 | 10000 |
| --- | --- | --- | --- | --- | --- | --- |
| Lumbrokinase sample | The fibrin plate | 8.37±0.17 | 11.13±0.18 | 12.44±0.32 | 13.65±0.28 | 14.68±0.15 |
|  | The fibrin-plasminogen plate | 16.32±0.10 | 18.60±0.07 | 19.04±0.05 | 19.94±0.13 | 20.29±0.28 |
| Lumbrokinase standard | The fibrin plate | 8.33±0.14 | 10.58±0.11 | 11.95±0.05 | 13.13±0.12 | 14.09±0.11 |
|  | The fibrin-plasminogen plate | 13.51±0.03 | 15.10±0.17 | 15.97±0.08 | 16.79±0.15 | 17.50±0.23 |

The current quality specification only measured its direct fibrinolytic activity by calibration curve and ignored the kinase activity. In our study, we observed that in the fibrin-plasminogen plate, the tendency of diameters of hydrolyzed clear zone between the sample and the standard was parallel, as shown in Fig 2. Furthermore, the parallel line method was employed to measure the total activity of lumbrokinase. As shown in table 3, the total potency calculation of all the five manufacturers ‘products were consistent with the method requirements, and three independent tests exhibited relatively good precision, suggesting that the parallel line method could be used to effectively measure total potency of lumbrokinase.

**Fig 2.**
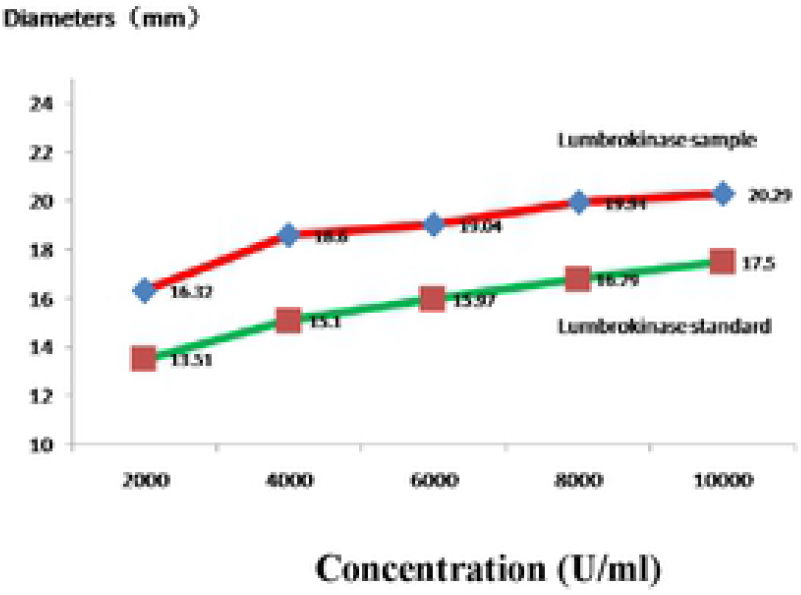
Diameter of fibrinolysis circle on the fibrin-plasminogen plate

**Table 3.** Total potency of lumbrokinase (%),calculated by parallel line method

| Total potency | 1# | 2# | 3# | Mean±SD | Confidence limit<br>(Mean) |
| --- | --- | --- | --- | --- | --- |
| Manufacturer A | 80.0 | 78.8 | 80.9 | 79.9±1.1 | 5.8 |
| ManufacturerB | 75.2 | 76.3 | 81.6 | 77.7±3.4 | 5.2 |
| ManufacturerC | 73.5 | 73.0 | 74.1 | 73.5±0.6 | 5.0 |
| ManufacturerD | 74.7 | 76.1 | 83.0 | 77.9±4.4 | 7.0 |
| ManufacturerE | 99.3 | 97.0 | 95.8 | 97.3±1.8 | 5.5 |

### Fibrin-zymography

The zymography was used to show the existence of lumbrokinase. Gel based zymography, combined with separation principle of polyacrylamide gel electrophoresis and enzyme-substrate reaction mechanism, could display the molecular weight of the active protein on the gel, and also could show the fibrinolytic band of lumbrokinase. According to the results of fibrin-zymography, there were six common active proteins in the products of the five manufacturers, as shown in Fig 3a. Besides, the products of one manufacturer had one more active protein band than others, and another one had two more bands. All the active bands were distributed in the range of 15~40KD.This suggested that lumbrokinase in the drugs was a group of fibrinolytic proteins. Subsequently, the consistency between different manufacturers was further evaluated, as shown in Fig 3b. Except for the little difference between two companies, the active ingredients and manufacturing process of all products of the five manufacturers were highly similar.

**Fig 3.**
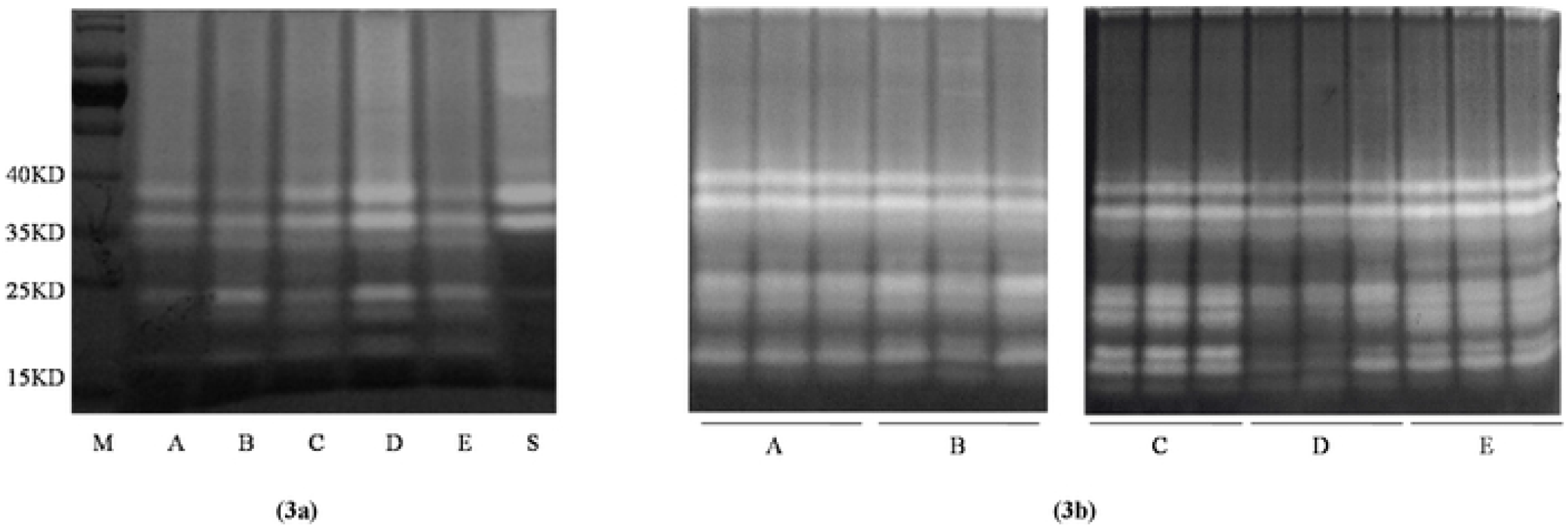
Fibrin-zymography reveals the active ingredients in the lumbrokinase drug substanecs. (3a)fibrin-zymography of lumbrokinase from five manufacturers and lumbrokinase standard. (3b) evaluation of consistency between different manufacturers, each manufacturer has three batches (M: protein marker; A~E: manufacturer A~E: S:lumbrokinase standard).

According to the results, lumbrokinase was a mixture of fibrinolytic components and exhibited different molecular masses in the polyacrylamide gel electrophoresis. Then, the species of earthworm were identified based on these results, and the method of fibrin-zymography was further verified.

In the specificity test, as shown in Fig 4a, inactive lumbrokinase and urokinase could not produce fibrinolytic lines. Defìbrase from snake showed two fibrinolytic lines, whereas they were obviously different from the lumbrokinase. The fibrinolytic lines between the *Eisenia foetida* and *Pheretima aspergillum* were overall different, revealing that fibrin-zymography could identify the active components containing the fibrinolytic protein and distinguishing *Eisenia foetida* from other earthworm species.

**Fig 4.**
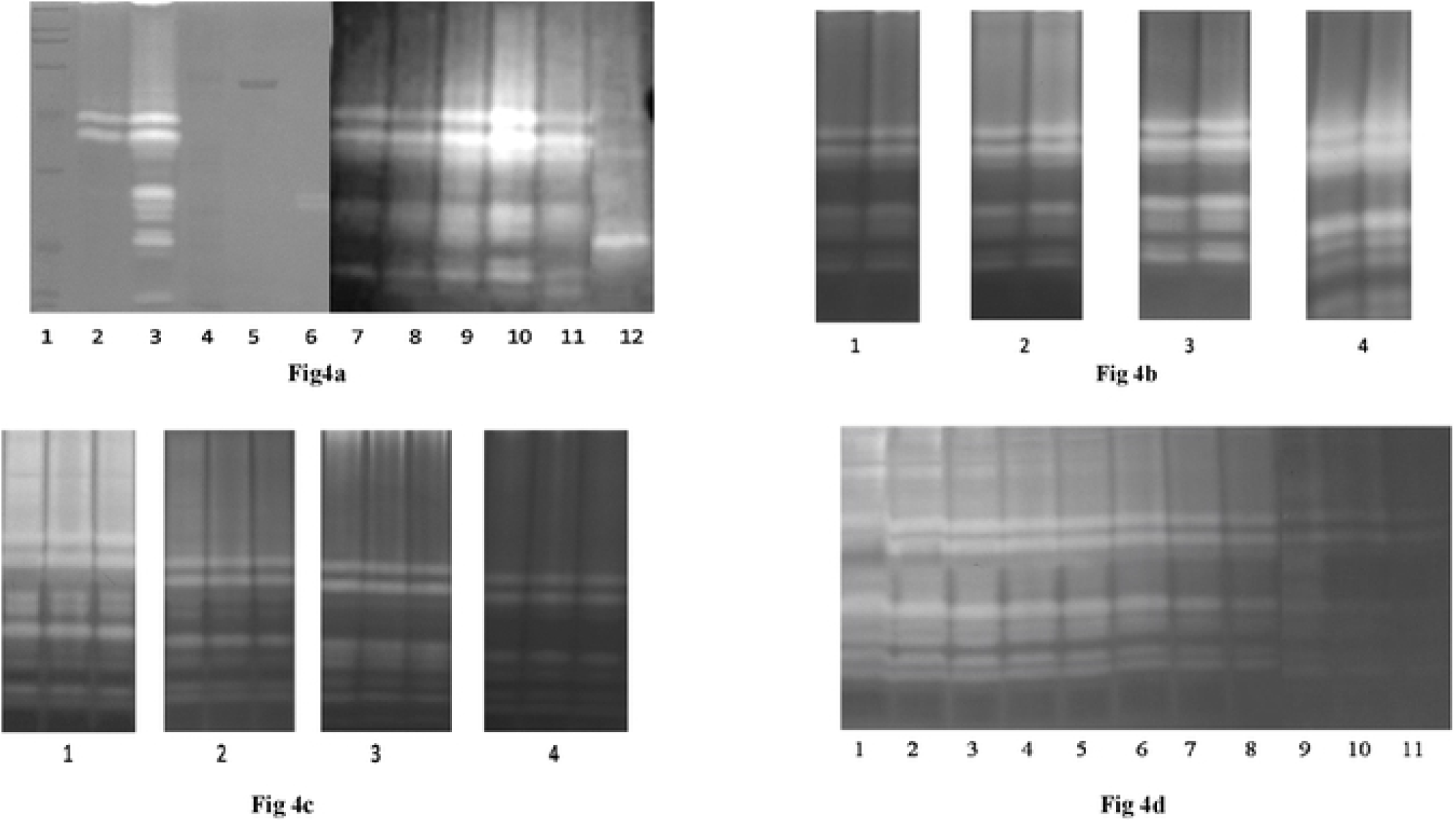
Verification Of fibrin-zymography. 4a specificity test: (1)marker. (2)lumbrokinase siandard:(3)lumbrokinase sample(4)inactive lumbrokinase sample:(5)urokinase:(6)defibrase(7-11)manufacturer A/B/C/D/E: (12)Pheretimaaspergillum: **4b incubation time**: (1) 10mins:(2)30mins:(3)60mins:(4) 120mins; **4c fibrinogen concentration 4d limit of quantitation**: (1) 0.25mg/ml: (2) 0.5mg/ml; (3) 1.0mg/ml: (4) 1.5mg/ml; **4d limit of quantitation**: (1-125μg:2-75μg;3-50μg:4-37.5μg:5-25μg:6-20μg.7-10μg;8-5μg:9-2.5μg.10-1.5μg.11-0.5μg)

In the robustness test, as shown in Fig 4b and 4c, the effects of fibrinogen concentration and incubation time on the clarity of fibrin-zymography were explored. The results showed that when the fibrinogen concentration ranged from 0.25to1.0mg/ml, and the incubation time ranged from 10 to 60 min. Accordingly, the fibrinolytic bands could be easy to identify.

In the test of range of quantitation, as shown in Fig 4d, a clear figure of fibrin-zymography could be generate when lumbrokinase concentration ranged from 0.5 to 37.5ug.

Based on the validation results, the method of fibrin-zymography satisfied the validation requirements, and it could be used as the identification method of *Eisenia foetida.*

### Proteins identification in lumbrokinase drug substances by electrospray ionization mass spectrometry(ESI-MS)

According to the results of fibrin plate experiments, lumbrokinase had two mechanisms of lysing fibrin, and we confirmed the presence of fibrinolytic protein in lumbrokinase using fibrin-zymography. Next, we identified the protein in lumbrokinase drug substances by LC-MS. The drug substances were treated with chymotrypsin. As shown in Table 4, fifteen fibrinolytic activity-related components were identified, and no other proteases were identified. 8-9 fibrinolytic proteins in each manufacturer were detected. Fibrinolytic protease 1, fibrinolytic protease P-III-1and protein ARSP1 existed in all products. The results of mass spectrometry suggested that lumbrokinase drug substances contained fibrinolytic and kinase components, which was consistent with the results of lumbrokinase activity measurement.

**Table 4.** Proteins identified by ESI-MS from five manufacturers.

| Manufacturer A | Manufacturer B | Manufacturer C | Manufacturer D | Manufacturer E |
| --- | --- | --- | --- | --- |
| Fibrinolyticprotease 1 (Q1ZZB7) | Fibrinolyticprotease 1 (Q1ZZB7) | Fibrinolyticprotease 1 (Q1ZZB7) | Fibrinolyticprotease 1 (Q1ZZB7) | Fibrinolyticprotease 1 (Q1ZZB7) |
| Fibrinolytic protease P-III-1(Q308Q8) | Fibrinolytic protease P-III-1(Q308Q8) | Fibrinolytic protease P-III-1(Q308Q8) | Fibrinolytic protease P-III-1(Q308Q8) | Fibrinolytic protease P-III-1(Q308Q8) |
| ARSP1(Q8I6N3) | ARSP1(Q8I6N3) | ARSP1(Q8I6N3) | ARSP1(Q8I6N3) | ARSP1(Q8I6N3) |
| Lumbrokinase-6 (Q6T373) | Fibrinolytic enzyme component A(Q8MX72) | Fibrinolytic enzyme large subunit(P81802) | Fibrinolytic enzyme component A(Q8MX72) | Fibrinolytic enzyme large subunit(P81802) |
| Fibrinolytic enzyme large subunt(P81802) | Lumbrokinase (A8ILP4) | lumbrokinase F238 (Q3HR18) | Fibrinolytic enzyme large subunit(P81802) | Lumbrokinase-6 (Q6T373) |
| Lumbrokinase F238 (Q3HR18) | Lumbrokinase (A8ILN1) | Lumbrokinase (A8ILP4) | Lumbrokinase-3 (Q6T376) | Lumbrokinase-5 (Q6T374) |
| Lumbrokinase (A8ILP4) | Lumbrokinase (A8ILP1) | Lumbrokinase (A8ILP1) | Lumbrokinase (A8ILP4) | Lumbrokinase-4 (Q6T375) |
| Lumbrokinase (A5HNU5) | Lumbrokinase (A5HNU5) | Lumbrokinase (A5HNU5) | Lumbrokinase (A8ILP1) | Lumbrokinase (A5HNU5) |
| Lumbrokinase (A8ILP1) |  | Lumbrokinase-6 (Q6T373) | Lumbrokinase (Q6DKQ2) | Lumbrokinase (A8ILN1) |

Subsequently, using the sequence alignment tool, the amino acid sequence of these proteins was compared with the known serine proteases. As shown in Fig 5, lumbrokinase shared the common features with the human trypsin, bovine trypsin, tPA(tissue-type plasminogen like activator) and uPA(urokinase-type plasminogen like activator)as their conserved catalytic triad, histidine(H), aspartate(D) and serine(S) were identical. The results implied that these active sites were highly conserved in these serine proteins, playing an important role in the activity of proteolysis.[13] Some lumbrokinases exhibited a highly similar loop and specific pocket (Asn-Asp-iLe-Ala-Leu-Leu), similar to tPA and uPA.[14–16] Besides, the amino acids, asp^507^, ser^532^ and gly^534^ in tPA, i.e. important substrate recognition sites for tPA and uPA, [17–19]also existed in several lumbrokinases. These proteins probably had the same activity with tPA or uPA.[20–23] The results suggested that some lumbrokinases might play a fibrinolytic activity, and the others might be able to lyse fibrin indirectly by activating the plasminogen to plasmin in the lumbrokinase related products.

**Fig 5.**
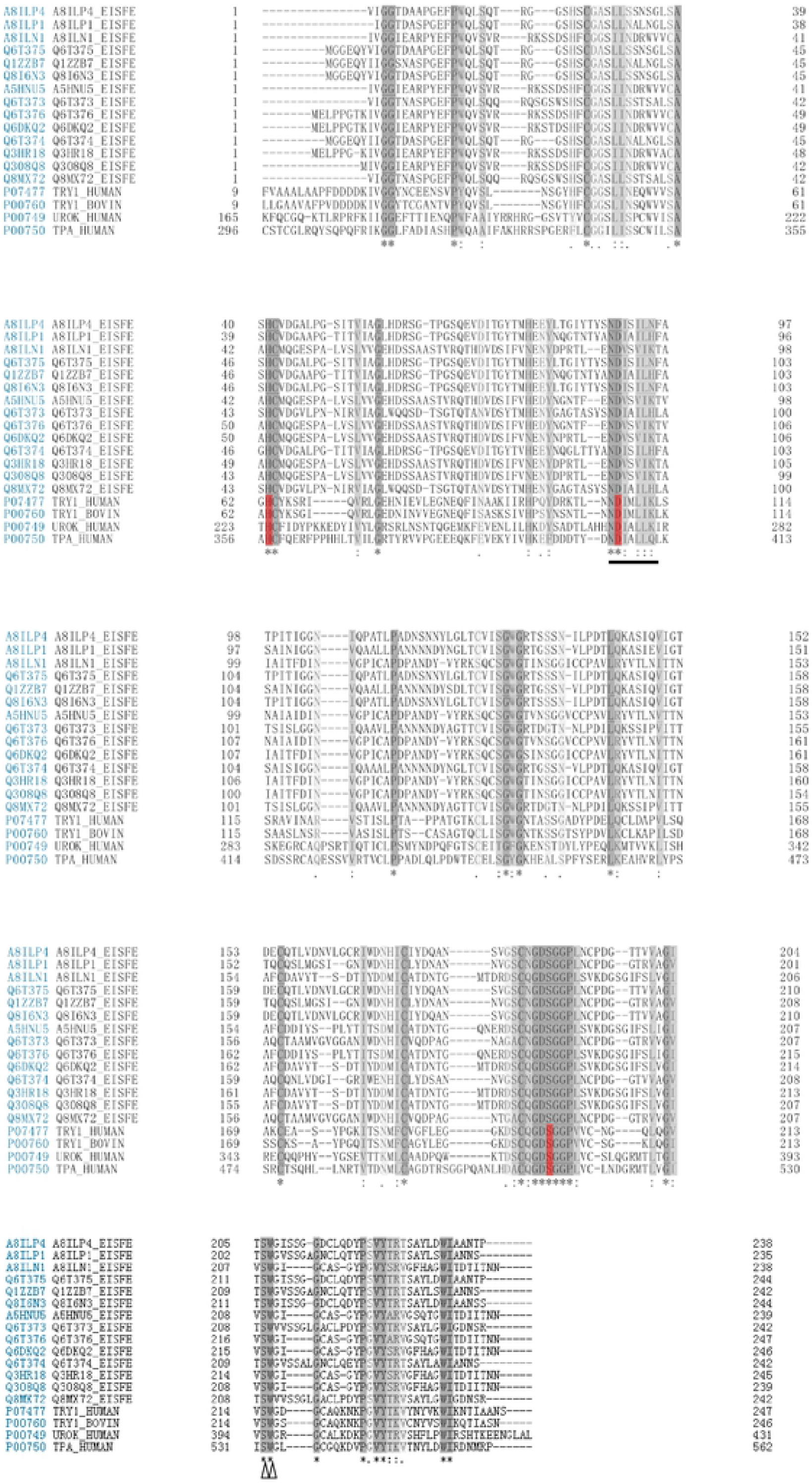
Sequence alignment of the identified proteins of E.fetida, human trypsin, bovine trypsin, human tPA and human uPA. The N terminal(or the C-terminal) part of The sequence was omitted The numberings shown were based on those proteins, respectively. Gaps, indicated by a dash, were introduced in the sequences to maximize the homology. The amino acid residues of the catalytic triad were red, substrate recognition sites for tPA and uPA was indicated by an triangle (Δ). The loop and specific pocket was underlined.

## Discussion

The fibrin-zymography confirmed the presence of active proteins in the lumbrokinase. The results of fibrin plate revealed that lumbrokinase had dual action mechanisms, which was further verified using the mass spectrometry. These results were consistent with the report that the protein property of these proteases, by activating the plasminogen into plasmin, both had the activity of degrading the fibrin directly and the activity of degrading the fibrin indirectly.[14] However, the current fibrin plate method used in the potency determination of lumbrokinase only measured its direct fibrinolytic activity with its kinase activity excluding. Accordingly, the current method should be optimized to measure its total activity. Our study showed that it was theoretically feasible to achieve the potency improvement of lumbrokinase using the parallel method.

It was reported that lumbrokinase(LK) was a group of fibrinolytic isozymes with molecular weights ranging from 25 to 32KD. [2, 24–26] These hydrolytic enzymes existed in different earthworm species, including *Lumbricus rubellus,[27, 28]Lumbricus bimastus[9]* and *Eisenia* fetida[29–31]. However, in the drug specification, identification of lumbrokinase based on the hemolysis test could not differentiate the proteases from different species. Fibrin-zymography based on the mechanisms of polyacrylamide gel electrophoresis could provide dimensional separations, helping to resolve the complex isozymes of lumbrokinase with relatively high sensitivity. In this study, fibrin-zymography was first introduced to identify the species of earthworm based on the hydrolytic bands on the gel and used to show the existence of active protein in the drugs and to distinguish the earthworm according to the distribution of bright bands on the gel.

Lumbrokinase was a group of serine proteins that exhibited highly similar physical and chemical properties. The amino acid sequence of each protein was highly homologous.[32] The contents of lysine (K) and spermine (R) were low in the protein sequence, thereby resulting in poor digestion of trypsin. Moreover, the isoelectric point of lumbrokinase was about 3-5, close to optimal pH of pepsin. Given this, we found the phenomena of protein precipitation in the process of treating lumbrokinase drug substances with pepsin. Thus, chymotrypsin was selected as the treating enzyme since the restriction sites, phenylalanine, tyrosine, tryptophan were widely distributed in lumbrokinase.

It is noteworthy that the database of *Eisenia foetida* contained very few types of proteins due to the lack of related research. Most of the proteins were fibrinolytic related. Thus, the results of mass spectrometry revealed that the drugs contained fibrinolytic proteins, and no other proteins were identified. Accordingly, a further study is required with the gradual update of the database.

Since the lumbrokinase was not isolated and purified from the earthworm in our study, which proteins showed the fibrinolytic activity, which one showed the kinase activity and which one showed both could not be concluded. However, our study laid a foundation for the pharmacological study of lumbrokinase. To identify each component in the enteric-coated capsules is our future work.

## Acknowledgments

We thank PhD Yinghua Xu of National Institute For Food and Drug Control (NIFDC, China) for providing technical support and editorial guidance and PhD Xia Zhang for editorial assistance.

